# A high-speed OLED monitor for precise stimulation in vision, eye-tracking, and EEG research

**DOI:** 10.1101/2024.09.13.612866

**Authors:** Olaf Dimigen, Arne Stein

## Abstract

The recent introduction of organic light-emitting diode (OLED) monitors with refresh rates of 240 Hz or more opens new possibilities for their use as precise stimulation devices in vision research, experimental psychology, and electrophysiology. These affordable high-speed monitors, targeted at video gamers, promise several advantages over the cathode ray tube (CRT) and liquid crystal display (LCD) monitors commonly used in these fields. Unlike LCDs, OLED displays have self-emitting pixels that can show true black, resulting in superior contrast ratios, a broad color gamut, and good viewing angles. More importantly, the latest gaming OLEDs promise excellent timing properties with minimal input lags and rapid transition times. However, OLED technology also has potential drawbacks, notably Auto-Brightness Limiting (ABL) behavior, where the local luminance of a stimulus can change with the number of currently illuminated pixels. This study characterized a 240 Hz OLED monitor, the ASUS PG27AQDM, in terms of its timing properties, spatial uniformity, viewing angles, warm-up times, and ABL behavior. We also compared its responses to those of CRTs and LCDs. Results confirm the monitor’s excellent temporal properties with CRT-like transition times (around 0.3 ms), wide viewing angles, and decent spatial uniformity. Additionally, we found that ABL could be prevented with appropriate settings. We illustrate the monitor’s benefits in two time-critical paradigms: Rapid “invisible” flicker stimulation and the gaze-contingent presentation of stimuli during eye movements. Our findings suggest that the newest gaming OLEDs are precise and cost-effective stimulation devices for visual experiments that have several key advantages over CRTs and LCDs.

---

Research on visual perception or cognitive psychology requires accurate displays with precise timing. Precise and consistent stimulus timing is also critical for the measurement of psychophysiological responses, for example those in the electroencephalogram (EEG). Due to their excellent temporal properties, cathode-ray tube monitors (CRTs) have long served as the gold standard in these fields, but CRTs in good working conditions are nowadays hard tofind. Also, while their pulsed emissions provide exact stimulus onsets, CRTs do not support the display of truly continuous luminance patterns (Elze, 2010b). They also suffer from poor spatial independence (e.g., Pelli, 1997), and their phosphor persistence can delay stimulus offsets (Brainard et al., 2002). In many laboratories, CRTs have now been replaced by liquid crystal displays (LCDs), which come with their own drawbacks. In LCDs, each subpixel contains liquid crystals that twist or untwist to pass through or block the light emitted by a backlight. Because this reorientation of the liquid crystals takes time, LCDs have comparatively sluggish response times that also depend on the content displayed during the previous frame (poor temporal independence). The use of a backlight means that LCDs cannot display true black, limiting their contrast. Furthermore, LCDs typically show a strong drop-offin luminance at wide viewing angles (e.g., Ghodrati et al., 2015; Zhang et al., 2018) and peripheral screen locations (e.g., Kawamoto et al., 2017). Since these issues are inherent to LCD technology, they also affect modern gaming LCDs (Zhang et al., 2018) and specialized LCDs for vision research or electroencephalography (Ghodrati et al., 2015).

## Potential advantages of organic light-emitting diode (OLED) monitors

An alternative display technology increasingly used in consumer products is the organic light-emitting diode panel, simply called “OLED” in the following. OLED screens use organic compounds that directly emit light when subjected to an electric current, which allows for precise control over each (sub)pixel’s output. The absence of a backlight also means that pixels can show true black, supporting practically infinite contrast ratios, increased viewing angles, and a wide color gamut. Most importantly, OLEDs promise excellent timing properties (Elze et al., 2013) with fast and temporally independent transition times.

The potential of OLEDs as stimulation devices for visual research was demonstrated already more than a decade ago (Cooper et al., 2013; Elze et al., 2013; Ito et al., 2013; Matsumoto et al., 2014). However, the panels characterized in these early studies were still technically limited in some ways; in particular, by supporting effective refresh rates of only 60 Hz. This limits their usefulness in paradigms requiringfine gradations in stimulus duration (Poth et al., 2018) such as visual masking, rapid serial visual presentation, multisensory integration, temporary order judgments, or other assessments of psychophysical thresholds.

In late 2023, a new generation of high-speed OLEDs entered the market. Aimed at video gamers, these monitors offer refresh rates of 240 Hz and models with 480 Hz have already been announced. These affordable consumer-grade OLEDs are advertised as having extremely short input lags (the latency from sending the GPU command to the onset of the monitor’s response) and fast transition times (the time from the start of the response until the desired color or gray level is reached). Together with the high image quality generally provided by OLED technology, this makes them interesting candidates for time-critical experiments on vision and cognition.

There are also drawbacks of OLED panels. One is that their organic compounds degrade faster than the inorganic materials used in CRTs or LCDs. In particular, screen regions where the same pattern persists for extended periods (e.g., taskbars, logos) face the risk of image retention from burn-in. Modern OLEDs include various functions to mitigate burn-in risk. These include the occasional shifting of the entire screen image by a few pixels (“pixel shifting”), the dimming of static screen regions, or “pixel cleaning” cycles, during which the pixels are rapidly turned on and off to remove residual charges. Because these features need to be deactivated for precise experimentation, it is unclear whether burn-in limits the monitor’s lifetime under laboratory conditions.

A second and more serious drawback of OLED monitors are luminance saturation effects and spatial dependencies caused by the panel’s “Auto Brightness Limiting” (ABL) mechanism. To manage power consumption and thermal output, OLEDs dynamically adjust the luminance of presented stimuli according to the current Average Picture Level (APL), the overall luminance output of all pixels. While the panel can display a small bright stimulus at maximum luminance, this becomes impossible with a large stimulus. With some OLED monitors, this ABL behavior can be experienced during everyday use where the display will noticeably dim whenever a bright window of the graphical user interface is maximized to fill the screen. In contrast to the anti burn-in features discussed above, ABL is an inherent technical limitation of OLED panels, although different panels will vary with regard to the APL at which this mechanism kicks in.

Obviously, any ABL behavior is highly problematic for precise experimentation. In fact, ABL-like luminance saturation effects were already identified as a major drawback in earlier tests of OLED monitors for vision research and medical diagnostics (Cooper et al., 2013; Elze et al., 2013; Ito et al., 2013). However, considerable technical progress has happened with consumer-grade OLEDs since. In particular, since current models support comparatively higher luminance levels, it should be possible to prevent ABL by operating them well below their peak brightness output (see also Cooper et al., 2013), where these monitors should still offer high contrasts.

### Current study

Goal of the present work was to evaluate the recently introduced high-speed gaming OLEDs for experimental research. For this, we characterized one promising candidate, the *ASUS ROG Swift OLED PG27AQDM*, with a focus on its temporal properties, which we also compared to those of CRT and LCD monitors available in our laboratory. In addition to conducting standard tests, we were especially interested whether luminance artifacts due to ABL could be prevented. Finally, we assessed the monitor’s timing performance in two exemplary time-critical paradigms, detailed in the following.

### Practical test: Fast flicker

As afirst practical test, we assessed the OLED monitor’s ability to faithfully reproduce fast flickering stimuli, where each luminance step is only shown for a single frame (4.17 ms).

In electrophysiological research,flickering stimuli have been commonly used to track visuospatial attention (Lalor et al., 2007; Morgan et al., 1996; Regan, 1966). Recent variants of this approach, the Rapid Invisible Frequency Tagging (RIFT) technique, measure the coherence between a high-frequency (50 to 86 Hz) contrast-modulated sinusoidal flicker sequence and neural responses in the magnetoencephalogram (Minarik et al., 2023; Seijdel et al., 2023) or EEG (Arora et al., 2024). A key advantage of RIFT is that the flicker remains barely visible^1^ (Minarik et al., 2023; Seijdel et al., 2023). This is attributed to the fast flicker rate and the more gradual contrast changes provided by the inclusion of intermediate gray levels within each flicker cycle. So far, RIFT stimulation has necessitated the use of an expensive high-speed projector refreshing at up to 1440 Hz (Minarik et al., 2023). Given their fast transition times, we tested whether a high-speed OLED can also faithfully present fast flicker sequences with intermediate gray levels. If so, this would suggest that RIFT-like stimulation may be feasible with off-the-shelf equipment.

### Practical test: Saccade-contingent stimulation

Another area of study that requires low stimulation latencies are eye-tracking paradigms in which the screen updates in near real-time with gaze position. For example, in one long-standing line of research, viewers see a preview of an object in parafoveal or peripheral vision (e.g., a word, a face, or a pattern of dots) before they make a saccadic eye movement towards it (McConkie & Rayner, 1975; McLaughlin, 1967; O’Regan & Lévy-Schoen, A, 1983; Rayner, 1975). To investigate the impact of the presaccadic preview on the following postsaccadic processing of the object at the center of gaze, these studies typically include an invalid preview condition where the object is exchanged, altered, or displaced during the saccade (Cox et al., 2005; e.g., Deubel et al., 1986; Herwig & Schneider, 2014; McConkie & Rayner, 1975; Rayner, 1975; C. Wolf & Schütz, 2015).

A major technical challenge with saccade-contingent paradigms is to time the display change to happen during the eye movement, while the viewer’s visual thresholds are elevated. This is less of an issue with large and therefore longer-lasting saccades, but challenging in tasks like reading (Rayner, 1975; Slattery et al., 2011), where saccades can be as short as 20 ms. Furthermore, some paradigms even require the presentation of multiple stimuli (e.g., Buonocore et al., 2020) or entire motion sequences (Schweitzer & Rolfs, 2021) during the saccade. Trials with mistimed display updates are not only lost for the analysis, but also cause oculomotor disruptions and conscious awareness of the change (Slattery et al., 2011).

Saccade-contingent stimulation has traditionally been performed with CRTs, although their phosphor persistence has led to some spurious findings (Bridgeman & Mayer, 1983; Jonides et al., 1983; W. Wolf & Deubel, 1993). More recently, high-speed projectors have been introduced for this purpose (Schweitzer & Rolfs, 2020; see also Richlan et al., 2013), but these systems are still prohibitively expensive for most researchers. The good temporal properties of modern gaming OLEDs should make them attractive for saccade-contingent experiments. Here we tested which change latencies can be achieved by linking the OLED with a 1000 Hz eye-tracker. Specifically, our goal was to present a stimulus exclusively during a saccade, but not before or after, in order to measure the brain-electric responses (not reported here) to these intrasaccadic stimuli.

## METHODS

### Display properties and settings

We tested two different units of the “ASUS ROG Swift OLED PG27AQDM” monitor, purchased on the open market. The monitor features a matte 27-inch (59.0 × 33.4 cm)

WOLED (white OLED) sample-and-hold display with a maximum refresh rate of 240 Hz. We will simply call this monitor “OLED” in the following, and “OLED-A” or “OLED-B” whenever we refer to a specific unit. In all tests, the monitor was controlled via the display port and operated at its native resolution of 2560 × 1440 pixels (see Table 1). Tests were conducted in the monitor’s standard dynamic range (SDR) mode and with its default “Racing Mode” activated. The monitor’sfirmware version was MCM104. Features to mitigate burn-in risks (“Screen Move”, “Pixel cleaning”, “Screen Saver”, “Adjust Logo Brightness”) were switched offin the on-screen display. If not stated otherwise, the OLED was tested with its “Uniform Brightness” (UB) feature activated. This mode is intended to avoid luminance changes due to ABL. For most tests, the monitor was used at its factory gamma setting of 2.2. Only for certain tests, in particular the 60 Hz flicker recording, it was linearized to a gamma of 1 using a Datacolor Spyder 4 colorimeter. The variable refresh rate feature was deactivated.

**Table 1.** Compared monitors.

| Abbrev. | Manufact. | Model | Panel type | Year | Refresh rate (Hz) (tested) | Refresh rate (Hz) (max.) | Diameter & Native resolution | Hardware settings Brightness / Contrast |
| --- | --- | --- | --- | --- | --- | --- | --- | --- |
| OLED | ASUS | PG27AQDM (Unit A & unit B) | OLED | 2023 | 240 | 240 | 27" 2560×1440 | 100% (260 cd/m<sup>2</sup>) / 80% |
| LCD-1 | ASUS | VG279Q | IPS | 2020 | 144 | 144 | 27" 1920×1080 | 45% (260 cd/m<sup>2</sup>) / 80% |
| LCD-2 | Iiyama | Prolite GS773HS | TN | 2013 | 100 | 120 | 27" 1920×1080 | 100% / 50% |
| CRT | Iiyama | MA203DT-D VisionMaster Pro 513 | CRT | 2003 | 120 | 180 | 22" (tested at 1024×786) | 100% / 100% |

### Stimulus presentation

Stimuli were presented using PsychToolbox-3 (PTB; Kleiner et al., 2007) under Matlab R2023b on a 64-bit Windows 10 Enterprise computer equipped with a mid-range GPU (AMD Radeon RX 6400). In all tests, a TTL trigger was sent from the stimulation computer’s native parallel port (using the “io64” Matlab utility) to the bioamplifier recording the photodiode signal immediately after the execution of the “flip” command in PTB.

The creators of PTB recommend Linux for precise stimulation. We therefore repeated some of the tests under Ubuntu Linux (version 22.04.4 LTS), which ran on the same stimulation computer and with the same Matlab version (R2023b). Since timing tests under Linux yielded similar results as those under Windows, we will only mention them in the context of the UB feature, which did not function properly under Linux.

### Recording apparatus

Luminance was measured using a Konica-Minolta LS-150 spot photometer with an aperture of 1°. If not stated otherwise, the photometer was mounted on a tripod at a distance of about 80 cm from the monitor. Luminance time courses were recorded with a Brain Products Photosensor photodiode and sampled at 8 kHz using an*eego*mylab amplifier (ANT Neuro) that recorded without a time constant (DC recording). High-speed videos of the monitor were taken with a Casio EXILIM EF-X1 camera at 1200 frames/s. All monitors warmed up for at least 20-30 min before testing. Monitor temperature at the screen surface was measured with a Sovarcate HS090E non-contact infrared thermometer.

### Reference monitors (CRT & LCD)

The OLED’s luminance responses were benchmarked against those of three LCD and CRT monitors available in our lab. An overview is provided in Table 1. First, we compared the OLEDs to a CRT monitor, the Iiyama MA203DT-D, called “CRT” in the following. This monitor is from a popular series of CRT screens (“Vision Master Pro” series; for details see Elze, 2010a, 2010b) and has often been used in time-critical studies on visual perception (e.g., Dimigen et al., 2012; Slattery et al., 2011; Wichmann et al., 2010).

Second, we compared the OLED to two LCDs of the same 27” diameter. The first, the ASUS VG279Q, called “LCD-1” in the following, is a fairly recent 144 Hz gaming monitor based on an in-plane switching (IPS) panel. The second, the Iiyama Prolite GS773HS, called “LCD-2” in the following, is an older 120 Hz model featuring a TN screen. All monitors were operated at their factory defaults for contrast and gamma (2.2) values. Where possible, their “Brightness” setting was adjusted so that peak luminance matched that of the OLED (i.e., about 260 cd/m^2^); see Table 1. The CRT, LCD-1, and LCD-2 monitors were connected via VGA, HDMI, and DVI connectors, respectively.

### Temporal properties

To obtain photodiode response profiles for all monitors, a full-white square (RGB: 255, 255, 255; size: 152 × 152 pixels) was presented either near the top (Figure 1A) or in the center (Figure 1B) of an otherwise black screen (RGB: 0, 0, 0) for two frames. This stimulus duration of two frames allowed us to observe both the onset (here: black-to-white) and offset response (here: white-to-black) as well as any between-cycle fluctuations. Depending on the monitor, the two frames corresponded to durations between 8.33 ms (at 240 Hz) and 16.67 ms (at 120 Hz). For most plots, luminance was normalized to a range of 0 to 1 (maximum), individually for each monitor. We focus on two latency measures, determined using Matlab scripts: The monitor’s rise time (or onset transition time) was defined as the interval from the stimulus reaching 10% until reaching 90% of its peak luminance (following Elze et al., 2013). Theflip-to-90% time was defined as the interval between the trigger marking the execution of the flip command to the moment at which luminance reached 90%.

**Figure 1.**
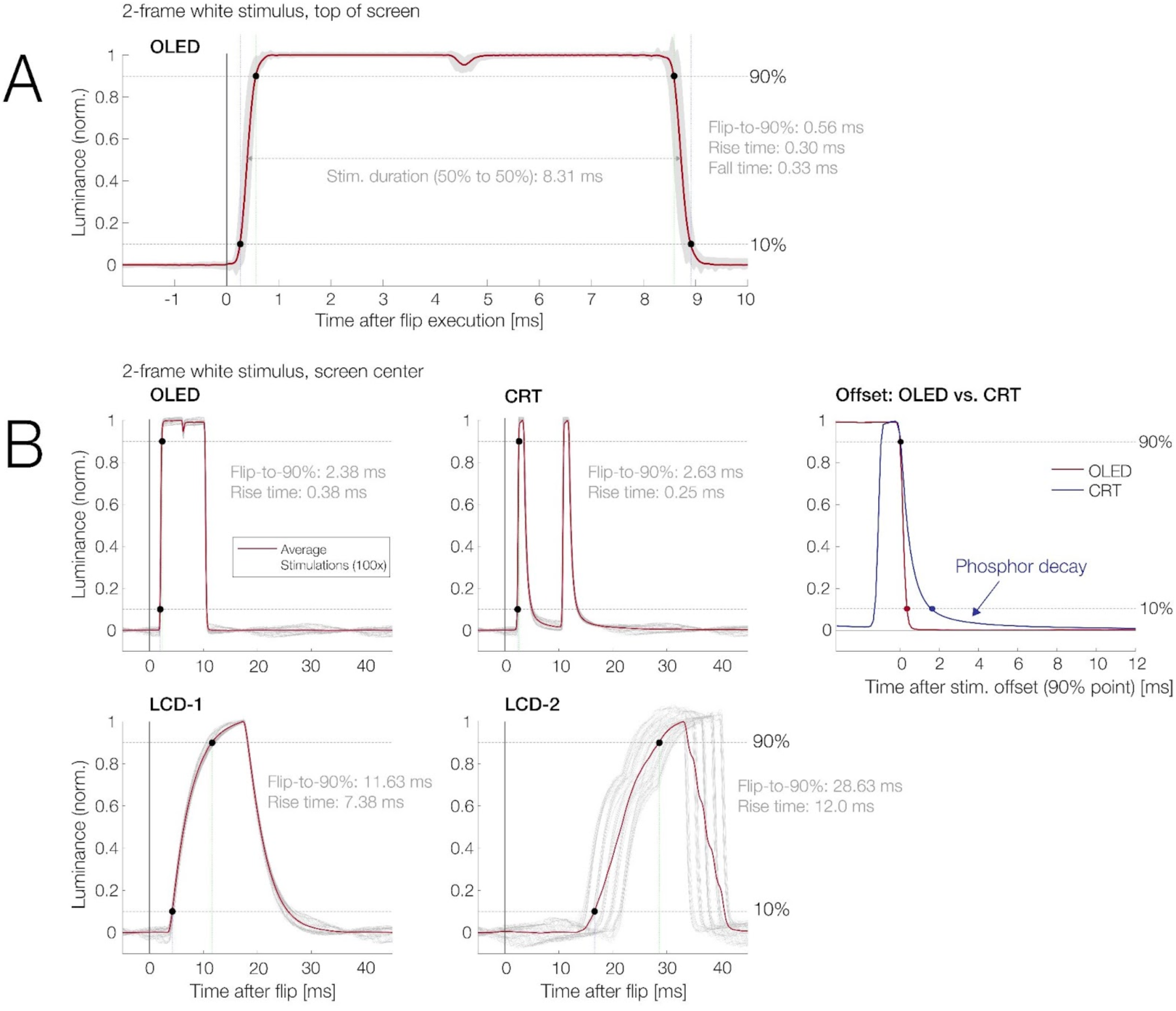
Luminance response to a white-on-black stimulus presented for two frames. In all plots, time zero marks the execution of theflip command and the red line is the average response to 100 presentations. Black dots mark the latencies at which the signal reaches 10% and 90%. (**A**) Enlarged view of the OLED’s luminance response. For this plot, responses were measured at the top of the screen. Gray shading indicates the region between the 2^nd^ and 98^th^ percentile of the 100 presentations. The slight dip in luminance between the two frames reflects OLEDflicker. (**B**) Comparison of the responses of different monitors to the same stimulus, but now presented in the screen center. Gray lines indicate individual responses. The OLED’s response latencies were comparable to those of a CRT monitor but the OLED did not show phosphor persistence at stimulus offset. Response latencies were far superior to those of the two tested LCDs.

### Spatial uniformity

A vision science monitor must exhibit spatial uniformity, meaning image properties should be as similar as possible at different locations. To assess luminance uniformity, we took spot photometer measurements from the center of 15 screen areas defined by a 5 × 3 rectangular grid (see Figure 2) while the OLED displayed a full-screen white stimulus (100% “Brightness”; UB feature activated). Measurements were conducted for both units of the OLED. We did not test uniformity for different colors.

**Figure 2.**
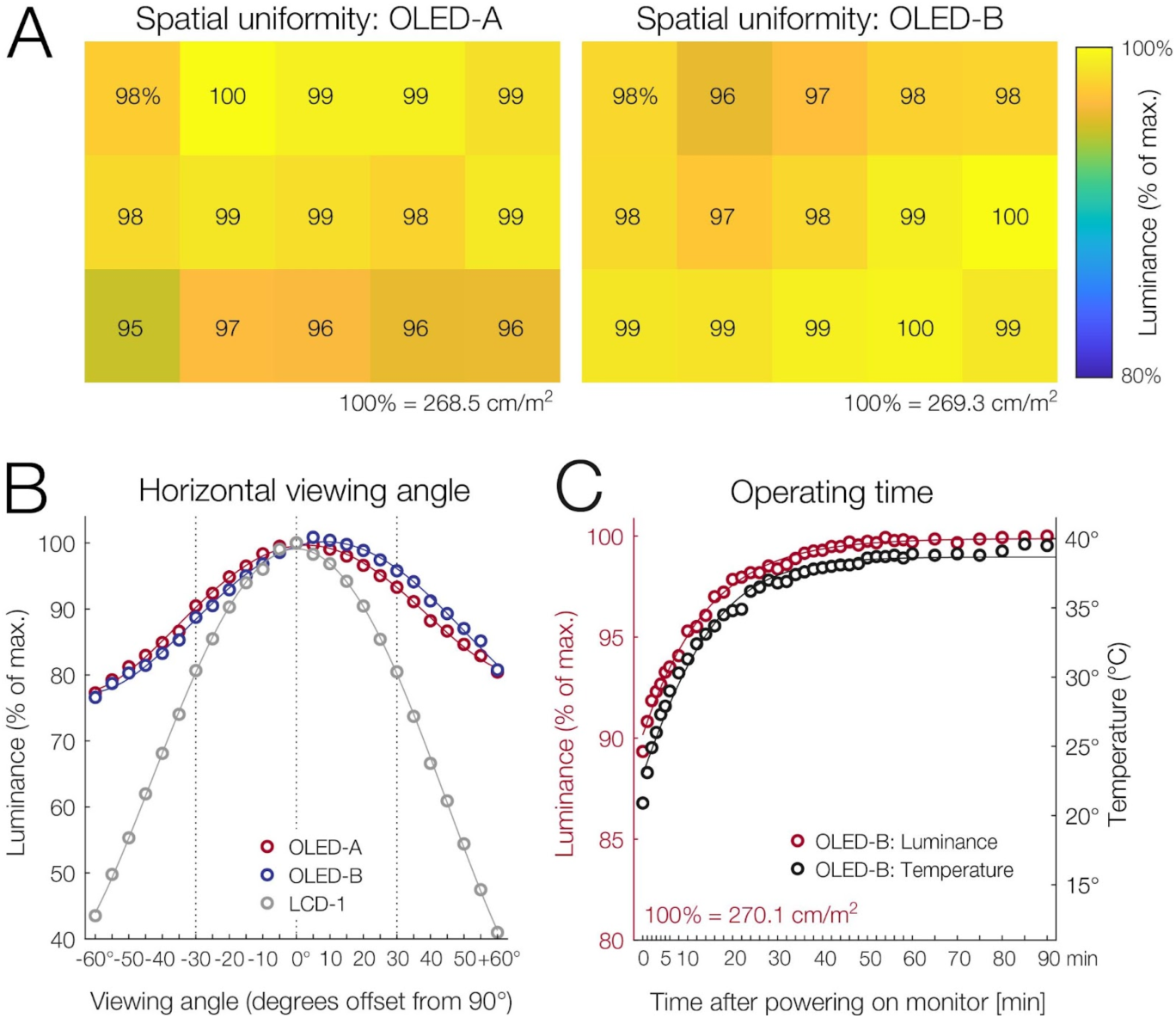
Luminance variations according to screen region, viewing angle, and operating temperature. During all measurements, the monitor showed a full-screen white stimulus. **(A)**Both tested OLED units showed decent luminance uniformity with a standard deviation of only 1.5% between the 15 measured screen regions (maximum: 5%).**(B)**Luminance of the OLED decreased by about 8% for viewing angle offsets of ±30° (dotted vertical lines). For comparison, the decrease was about 20% for LCD-1, a monitor with an IPS panel by ASUS.**(C)**Luminance increased while the monitor warmed up and reached an asymptotic luminance of 99% after 36 min.

### Viewing angle

To measure luminance as a function of horizontal viewing angle, photometer readings were taken at the screen center with the monitor rotated to different angular offsets (azimuths) spanning −60° to 60°. For this test, the OLED monitor again displayed a full-white stimulus and the test was conducted with both units. For comparison, viewing angle measurements were also taken for LCD-1. We did not test for shifts in color as a function of viewing angle.

### Operating time

We assessed to what extent the OLED’s luminance depends on its operating temperature and how long the monitor takes to reach its asymptotic luminance. To ensure accuracy, testing was conducted only after the monitor had cooled to room temperature (19.8°C) overnight. After turning on the monitor, spot photometer and infrared thermometer readings were taken at regular intervals (initially every minute, later every 2 min., then every 5 min.) from the screen center, while it continuously displayed a full-screen white background (at 100% “Brightness”; UB feature activated). To visualize the relationship, logistic functions were fitted to the temperature and luminance data (Figure 2).

### Auto Brightness Limiting

We used a common procedure to assess the monitor’s ABL behavior and to test the monitor’s UB feature that is intended to prevent luminance changes as a function of the APL. To manipulate the APL, a full-white rectangle was presented in the center of an otherwise black screen (RGB: 0, 0, 0). In subsequent measurements, the size of the rectangle was iteratively increased so that its area subtended 1%, 2%, 5%, 10%, 25%, 50%, 75%, or a full 100% of the screen area. Each APL level was displayed for about 10 seconds and a photometer reading was taken from the screen center, both immediately after stimulus onset (initial luminance) and about 3-5 s later (sustained luminance). The test was conducted both without and with the UB feature activated. Additionally, to capture a phenomenon that we will refer to as “dynamic dimming” below, we repeated the test with time-resolved photodiode recordings.

### Practical test: High-frequency flicker stimulation

We tested whether the response times of the OLED support the faithful presentation of fast flicker time series, similar to those used in the RIFT paradigm. Utilizing the 240 Hz refresh rate, we presented a circular flickering patch (a disc with tapered edges) on a gray (50%) background at a frequency of 60 Hz. To reduce visibility of the flicker, the time series included two intermediate gray states within each four-frame cycle (i.e., patch luminance of 0%, 50%, 100%, 50%, …). The key question was whether the OLED could reach the luminance values with high fidelity within each 4.17 ms frame.

### Practical test: Intra-saccadic stimulation

To illustrate the advantages of the monitor in a time-critical scenario, we implemented a gaze-contingent display change paradigm with intrasaccadic stimulation. Goal of this simple experiment was to present a stimulus exclusively for the duration of a brief saccadic eye movement.

The experiment is illustrated in Figure 6A. Each trial started with a whitefixation cross displayed on the left side (−10° eccentricity) of an otherwise uniform gray screen. Once a stablefixation on the cross was registered, a second white fixation cross – the saccade target – appeared at +10° eccentricity, thereby triggering the participant to execute a 20° rightward saccade. Only during the saccade, once the movement was detected online in the eye-tracking data (see details below), a circular grating of varying orientations (diameter: 16°, spatial frequency: 0.15 cycles-per-degree) appeared in the screen center. The stimulus offset of the grating was initiated once the gaze crossed an invisible vertical boundary placed 4° (or 210 pixels) to the left of the saccade target. Participants then judged the grating’s orientation with a button press.

A desktop-mounted Eyelink 1000 video-based eye-tracker (SR Research), controlled by the Eyelink extension for PTB (Cornelissen et al., 2002), recorded gaze position from the left eye at 1000 Hz. The left eye was chosen because it is the adducting eye for a rightward saccade, which shows smaller post-saccadic overshoots. To reduce processing delays, filters were deactivated for gaze samples streamed online to the stimulus computer via the Ethernet link. For eye movement data stored offline, the heuristic filter level was set to 1, meaning that the filter was applied across pairs of subsequent samples.

The photodiode was placed in the screen center, at a position that always became illuminated (turned white) when a grating was displayed. Data was recorded from one trained observer (one of the authors) who performed 200 trials, sitting at a distance of 62 cm from the screen. For eye comfort, the monitor was operated at a Brightness setting of 40% for this experiment (UB feature activated). For all critical events within each trial (defined in the next paragraph), a parallel port trigger was sent from the stimulus PC via a looped-through cable to both the eye-tracker and the photodiode’s bioamplifier. Offline, the photodiode data (downsampled to 2 kHz) was synchronized with the eye-tracking data using the EYE-EEG toolbox for EEGLAB (Dimigen et al., 2011). Synchronization was based on >1000 shared trigger pulses sent to both Eyelink and the amplifier. The average synchronization error between both time series – computed based on trigger alignment in both recordings after synchronization – was 0.38 ms.

For each trial, we assessed the latency of six critical events. The first was the onset of the saccade (*t*_saccOn_), as detected offline after the experiment. Offline saccade detection was performed with the widely used Engbert & Kliegl (2003) algorithm with typical parameters (adaptive velocity threshold: λ=6 median-based SDs, min. saccade duration = 8 ms).

The second latency, t_saccDetect_, defined the moment that the saccade was detected online using the method of Schweitzer & Rolfs (2020). Their algorithm, available as compiled C-code (https://github.com/richardschweitzer/OnlineSaccadeDetection) is a variant of the Engbert & Kliegl (2003) algorithm optimized for online use. Beginning with the presentation of the saccade target, new gaze position samples were retrieved every millisecond from the Eyelink and passed to the function together with all previous samples from the trial. A saccade was detected online once eye velocity exceededλ=9 median-based SDs for two consecutive samples (2 ms) and the movement had an angular orientation to the right (±45°).

The third latency,*t* _flip_, marks the moment the flip command was executed in PTB, which exchanges the GPU’s front and back buffers. The fourth latency,*t* _stimOn_, indicates the actual stimulus onset in the screen center, as determined offline in Matlab. Like before, it was defined as the moment when stimulus luminance exceeded 90% of its sustained peak luminance. The fifth latency,*t* _StimOff_, marks the stimulus offset, defined as the latency at which luminance dropped again below 10% of the peak level. The final latency was the end of the saccade,*t* _SaccOff_, as detected by the offline algorithm.

We focus here on several critical intervals defined by these events:

(1)*SaccOntoSaccDetect*
(2)*SaccDetecttoFlip*
(3)*FliptoStimOn*

The latter two intervals depend strongly on the monitor’s properties: Interval (2) is largely determined by monitor refresh rate because it influences when the nextflip can happen. Interval (3) is determined both by refresh rate (because it determines how long it takes for the update to arrive at the screen center) but also by the monitor’s rise time.

These intervals can also be combined into (4) the*display change latency*(as commonly reported, for example, in reading research), which is the sum of (2+3). Finally, the (5)*total latency*was defined as the entire interval from saccade onset to stimulus onset (1+2+3).

## RESULTS

### Temporal properties

Figure 1A shows the average response of the OLED monitor to 100 presentations of a white stimulus on a black background. The stimulus was shown for two frames, corresponding to a nominal stimulus duration of 8.33 ms (2 × 4.167 ms). For Figure 1A, the stimulus was presented near the top of the screen (about 70 pixels from its upper edge). Time zero marks the timing of a trigger sent at the execution of the “flip” command.

Both onsets (black-to-white transitions; rise time) and offsets (white-to-black transitions; fall time) were extremely fast: They lasted 0.3 ms (rise) and 0.33 ms (fall) and were almost perfectly symmetrical. Relative to the flip, the stimulus reached 90% of its peak luminance within about half a millisecond (0.56 ms). In-between the two display frames, there was a dip in luminance of about 5%. This dip reflects an OLED flicker at the 240 Hz refresh rate (see next section). Otherwise, the OLED’s luminance response was close to a square wave (Elze et al., 2013; Matsumoto et al., 2014). The horizontal arrow in Figure 1A highlights the interval between the 50% luminance points, which were 8.31 ms apart. The monitor’s duty cycle therefore closely matched the nominal stimulus duration of 8.33 ms (two frames of 4.17 ms each).

Figure 1B compares the OLED’s luminance response to that of selected CRT and LCD monitors. For the results in Figure 1B, the white stimulus and the photodiode were placed in a more common location for all monitors, the screen center. Since the OLED updates from top to bottom, this adds some delay relative to the flip latency. As Figure 1B shows, the rise times of the OLED (in this test about 0.38 ms) were only slightly longer than those of the CRT monitor running at 120 Hz (0.25 ms). Notably, however, the mean flip-to-90% latency was even shorter for the OLED (2.38 ms) than the CRT (2.63 ms).

In contrast to the OLED, the CRT showed asymmetric on- and offset responses with a slower fall time than rise time. The rightmost panel in Figure 1 compares the offset responses in detail. For this panel, responses were temporally aligned to stimulus offset (i.e., the point when luminance falls below 90%). Whereas the OLED’s luminance dropped to zero (dark screen) within half a millisecond, the CRT showed a more gradual offset response due to phosphor persistence. After the CRT’s luminance had fallen below 10%, it took another 10 ms to decay to 1% luminance and another >30 ms to decay entirely (to 0%).

The bottom panels in Figure 1B show the responses for the two tested LCDs. The results exemplify the problems seen with many LCD screens (Ghodrati et al., 2015): Sluggish responses and asymmetrical on- vs. offset profiles. The newer LCD-1 screen had a rise time of 7.38 ms and reached 90% luminance only 11.63 ms after the flip. The older LCD-2 had even slower transitions and also showed synchronization issues with the flip command (as visible in the single-trial response in Figure 1B).

### OLED flicker

As visible in Figures 1A and 4C, the OLED exhibits a typical flicker at its refresh rate of 240 Hz. Flicker in OLEDs typically results from brief voltage changes at the pixel’s transistor gates which control the light output during each refresh cycle. This subtly alters the pixel’s brightness for a brief moment. While the exact amplitude of this 240 Hz flicker varied between our tests and according to the displayed luminance level, it was generally small. For example, in the data shown in Figure 4C (full-white stimulus, 260 cd/m^2^), its peak-to-trough amplitude was around 4.6%, or 12 cd/m^2^.

### Spatial uniformity

Figure 2A illustrates the luminance uniformity for two different units of the OLED. For a full-white stimulus, displayed at an average luminance of around 260 cd/m^2^, luminance variations never exceeded 5.5% and were mostly much smaller. The standard deviation across 15 measured screen locations was 1.5% (or 4.03 cd/mm^2^) for OLED-A and 1.1% (or 2.96 cd/mm^2^) for OLED-B. Across our two tested units, the pattern of inhomogeneities was unsystematic, i.e., there was no clear luminance peak in the screen center as sometimes seen with CRTs and LCDs (Kawamoto et al., 2017; Zhang et al., 2018).

### Viewing angle

Figure 2B depicts luminance as a function of viewing angle for the two OLED units. At viewing angle offsets (azimuth) of ±30°, luminance dropped by about 8% to a value of 92.07% (value averaged across both units and across positive and negative offsets). In comparison, for LCD-1, a fairly recent ASUS gaming monitor using an IPS panel, the drop-offwas almost 20% (to 80.5%, averaged across positive and negative angular offsets). At extreme offsets of ±60°, luminance dropped by 21.2% (to 78.8%) for the OLEDs and by 57.8% (to 42.2%) for LCD-1. For both OLED units tested, there was a slight asymmetry for negative versus positive angular offsets, which was not present for LCD-1.

### True Black

While showing a full-screen black stimulus, the OLED did not produce light emissions measurable with our photometer. Subjectively, the screen remained invisible after 15 min of dark adaptation in a totally dark room. A long exposure photograph (tripod-mounted Sony Alpha 6000 camera, Samyang 12 mm lens set to an F2.0 aperture, exposure time: 30 s at ISO 20,000) also provided no evidence of light emissions.

### Operating time

After switching on the monitor, luminance increased steadily over 90 min from an initial level at room temperature of 91.6% (or 241.3 cd/mm^2^) to the full 100% once the monitor had warmed up to 39.5 °C (Figure 2C). However, an asymptotic luminance level of 99% (267.8 cd/mm^2^) was already reached after 36 min of operation (at 37.6 °C).

### Auto-Brightness Limiting behavior

The Uniform Brightness feature is intended to eliminate local luminance changes due to changes in the Average Picture Level. Figure 3 shows the luminance of a white stimulus as a function of its size (i.e., the APL) both without (panel A) and with (panel B) the UB feature activated. Without the UB feature (Figure 3A), stimulus luminance changed drastically with the APL whenever the monitor was operated at “Brightness” settings above 40%. The 40% setting corresponds to a luminance of about 140 cd/m^2^. Importantly, however, at settings of 40% or less, no ABL occurred. This suggests that ABL can be reliably prevented by operating the monitor at moderate brightness settings.

**Figure 3.**
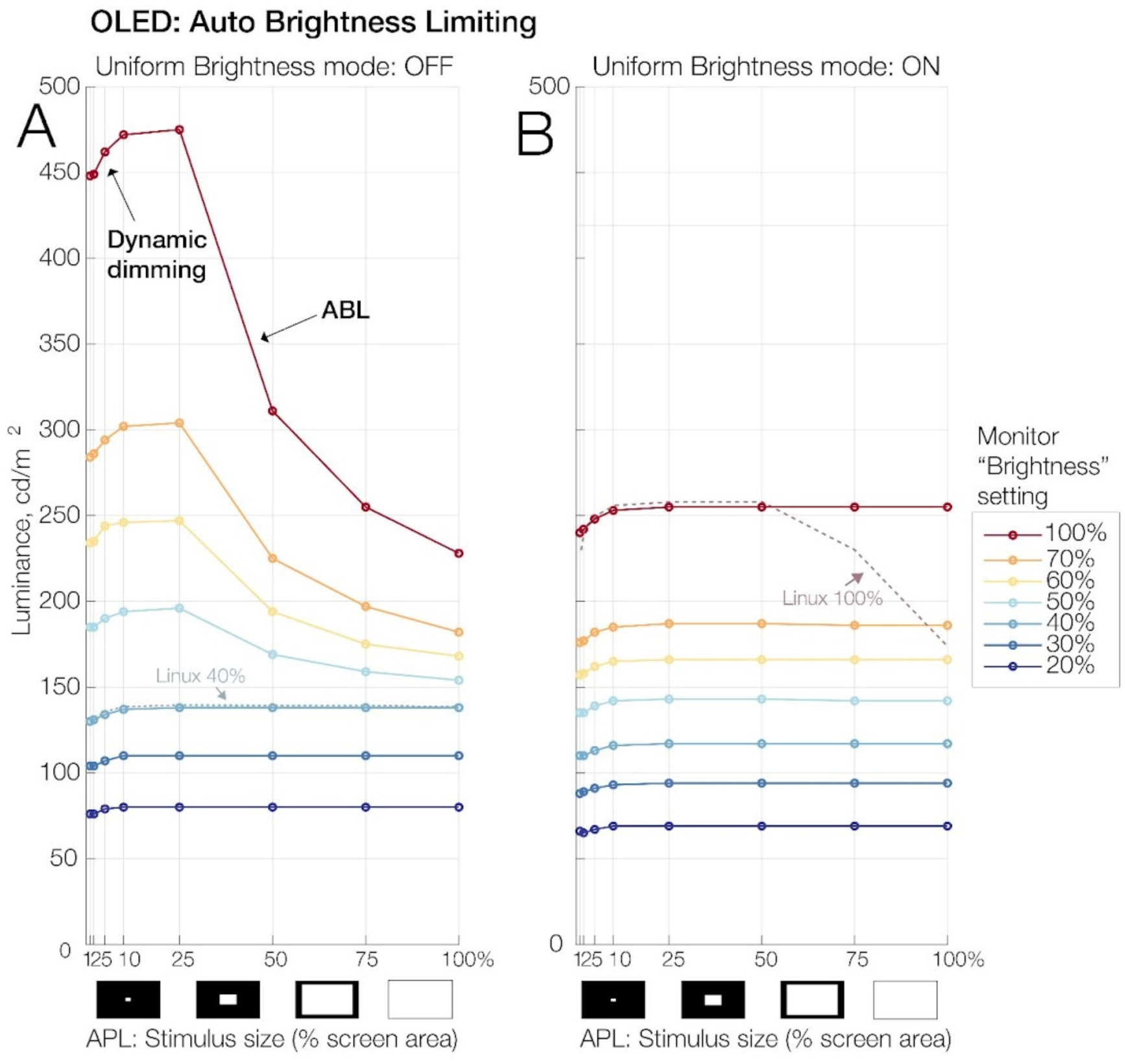
Auto Brightness Limiting (ABL) behavior of the OLED as a function of the Average Picture Level (APL), the monitor’s “Brightness” setting, and the activation of the “Uniform Brightness” feature. The abscissa indicates the area (in % of total screen)filled by a full white stimulus presented on an otherwise black background. This is equivalent to the APL, the monitor’s overall luminance output. As can be seen, sustained stimulus luminance decreased drastically with increasing APL, a typical behavior of OLEDs. Importantly, however, ABL could be reliably prevented by operating the monitor at “Brightness” settings of 40% or less (panel A) or alternatively, by activating its “Uniform Brightness” feature (panel B) which prevented ABL at all Brightness settings. The slight decrease of sustained luminance for small bright stimuli (10% area or less) visible in all plots is caused by “dynamic dimming”, detailed in Figure 4. Notably, the Uniform Brightness feature did not work properly under Linux as exemplified here for a Brightness setting of 100% (panel B, dotted line). However, ABL could also be prevented under Linux by running the monitor at or below 40% Brightness (panel A, dotted line).

Figure 3B shows the results from the same test with the Uniform Brightness feature activated. At any given Brightness setting, the UB feature slightly decreased the monitor’s luminance. More importantly, however, it prevented dimming due to ABL at all Brightness settings, meaning that the monitor’s brightness no longer changed according to the APL. With this setting, Brightness could be safely increased to a setting 100% (corresponding to about 260 cd/m^2^) without any measurable ABL behavior.

We repeated this test also under Ubuntu Linux. To our surprise, we found that the UB feature did not work properly under Linux. This is exemplified in Figure 3B for the condition with 100% Brightness (dotted line). This means that Linux users need to operate the monitor at Brightness settings of ≤40% (see dotted line in Figure 3A) to prevent ABL.

### Dynamic dimming

We observed an unexpected dimming behavior that happened exclusively for relatively small and bright stimuli (Figure 4). Specifically, we found that the luminance of full-white stimuli subtending <10% of the screen area of an otherwise dark screen decreased in luminance over the first 500 ms after stimulus onset. The magnitude of this decrease was up to 5% in the photometer data (Figures 4A and 4B). In the photodiode data, shown in Figure 4C, the decrease was about 2% for the tested stimulus, which subtended 2% of the screen area. After 500 ms of a linear decrease, the luminance plateaued. This behavior, which we call “dynamic dimming”, was not observed for larger stimuli (≥25% of screen area). Also, unlike ABL, which was effectively prevented by activation of the UB feature, this dynamic dimming occurred regardless of it.

**Figure 4.**
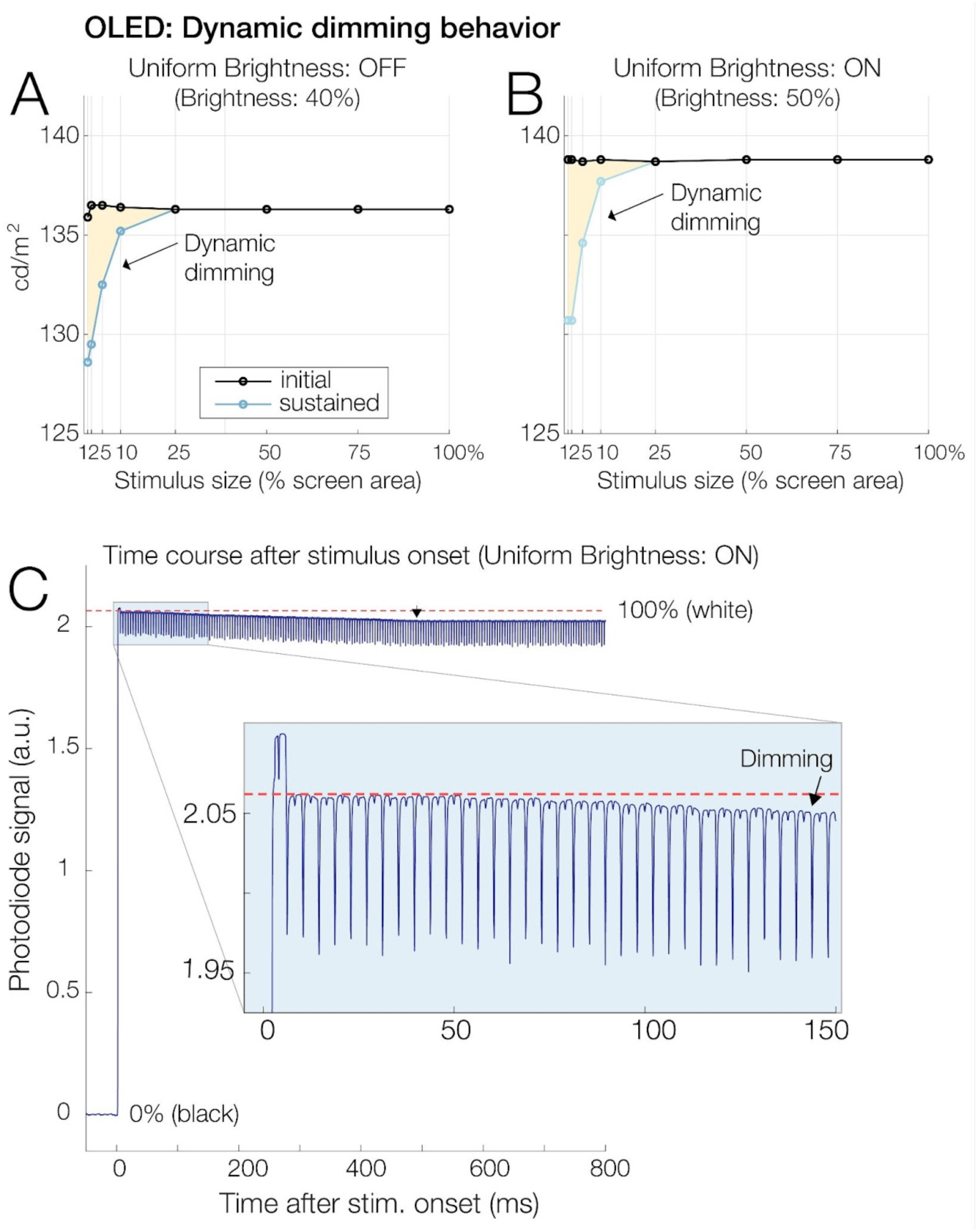
Dynamic dimming of small bright stimuli. (**A**) Spot photometer readings for a white (100%) stimulus of varying size presented on a black background. Readings were taken immediately after stimulus onset (“initial”, black line) and 3-5 s later (“sustained”, blue line). Interestingly, small (≤10% of screen area) but not large stimuli dynamically decreased in luminance by about 2-5%. This happened even at low “Brightness” settings at which ABL (see Figure 3) did not occur anymore. (**B**) In contrast to ABL, this dynamic dimming was seen regardless of the “Uniform Brightness” feature. (**C**) Photodiode time course of dynamic dimming (average of 100 stimulations) for a small (2% of screen area) white stimulus. Luminance decreases over the first ∼500 ms, but then plateaus.

### High-frequency flicker stimulation

Figure 5 shows an exemplary 40 s of aflickering patch presented at 60 Hz with four frames per cycle. Due to the rapid transition times of the OLED (of only 0.3-0.4 ms, see Figure 1) each of the 4.17 ms long flicker states could be faithfully presented at the intended duration. Relative to the intermediate gray value of 50%, there was some asymmetry in response amplitude for the bright states (100%) as compared to the dark states (0%); this may have been due to an imperfect gamma calibration of the monitor.

**Figure 5.**
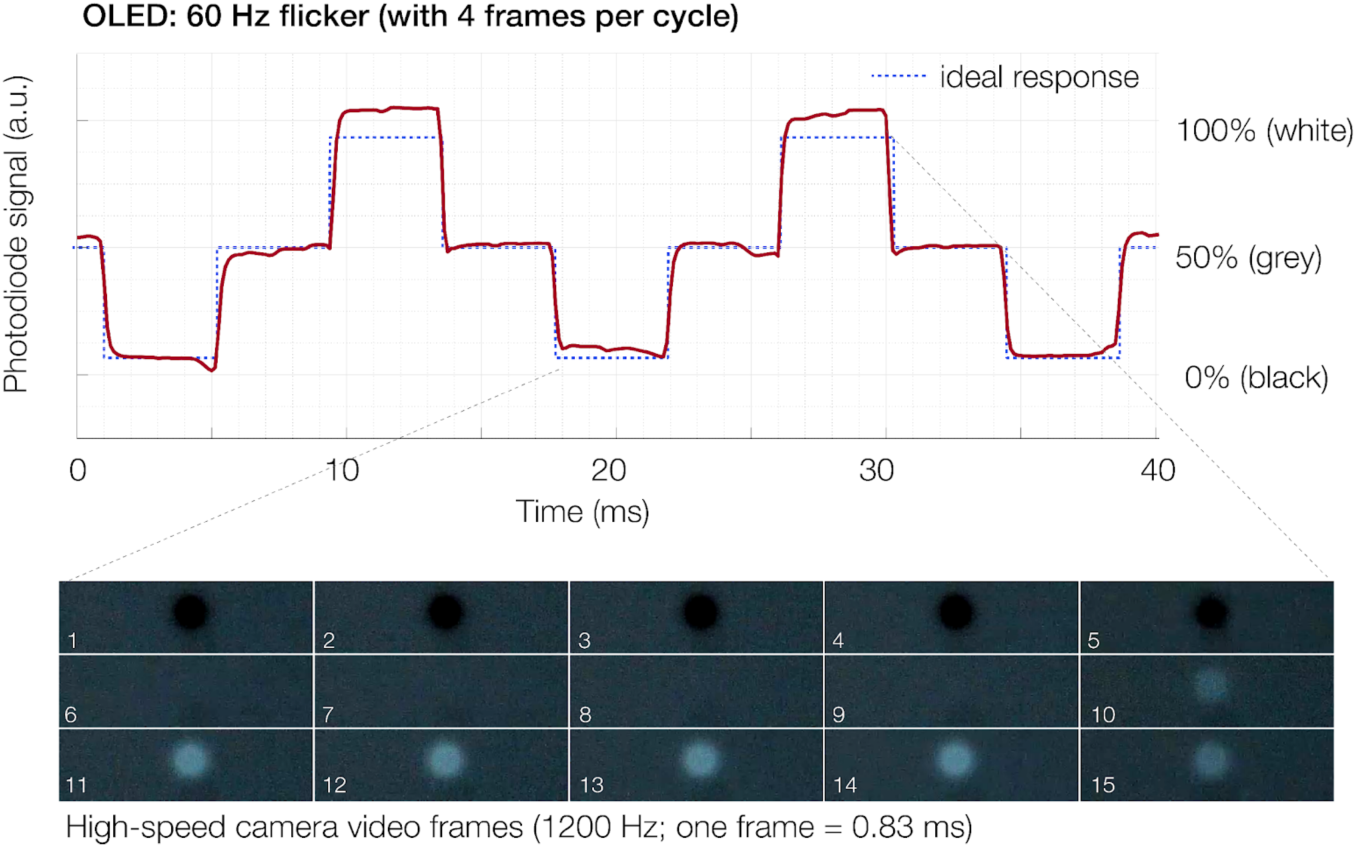
Upper panel: Luminance response for a circular contrast-modulated patch flickering at 60 Hz (background: 50%; luminance of patch: 0%, 50%, 100%, 50% etc.) with an intermediate gray level to reduce theflicker’s visibility, as in Rapid invisible frequency tagging (RIFT). The dotted blue line indicates the theoretical ideal response. Lower panel: Frames from a high-speed video of the flicker sequence. Each video frame lasts 0.83 ms.

### Intra-saccadic stimulation

The online saccade detection led to some premature display updates before saccade onset (false alarms; 9.5% of trials), caused by microsaccades or unstable tracking of the eye. Trials with false alarms or other issues (e.g., eye blinks, saccades that were too short) were removed for the current analysis (total: 17% of all trials). Data of the remaining trials of one observer is visualized in Figure 6 (using the Violinplot-Matlab package; Bechthold, 2016). As can be seen, it was routinely possible to change the screen well before the saccade reached its peak velocity. The mean durations of the critical intervals were as follows:

(1)*SaccOn*to*SaccDetect*:5.28ms (SD:1.84ms)
(2)*SaccDetect*to*Flip*:4.69ms (SD: 1.19ms)—depends on monitor
(3)*Flip*to*StimOn*:2.92ms (SD:0.29ms)—depends on monitor

Relative to the online detection of the saccade (*SaccDetect*), the display was updated on average with a*display change latency*of 7.61 ms (SD: 1.18 ms). Relative to the saccade’s actual onset, as detected offline with more sensitive thresholds, the stimulus onset occurred with a*total latency*of 12.89 ms (SD: 2.25 ms). The average saccade duration was 60.36 ms; of this, it was possible to show the stimulus on average for 39.33 ms, or 65.16% of the saccade. Only in rare cases (5 of 200 trials), the stimulus was switched off too late, after the saccade had landed.

**Figure 6.**
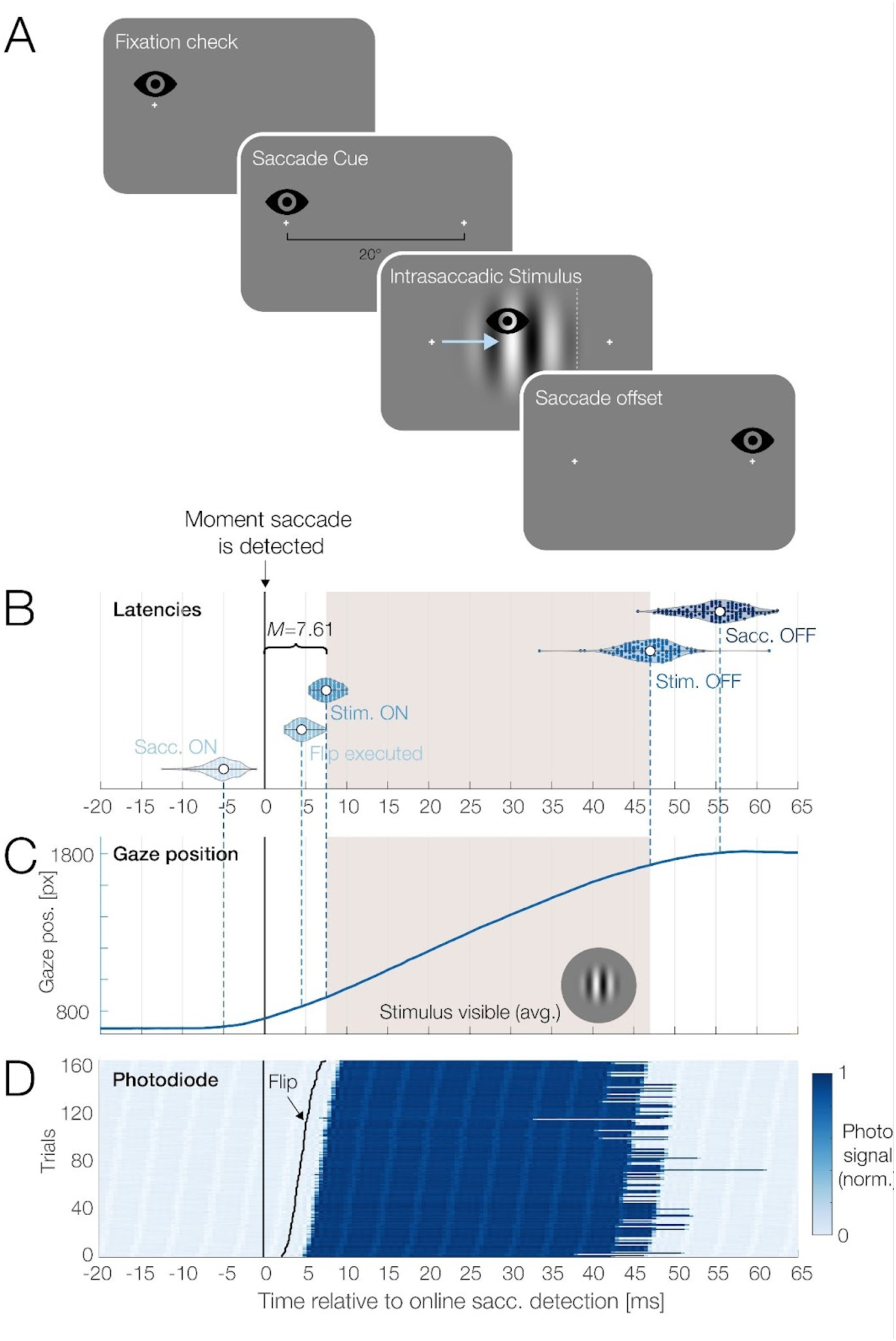
Practical test in an experiment with gaze-contingent intrasaccadic stimulation. Goal is to show a grating stimulus only for the duration of a saccade, but not before or after. **(A)**Following a fixation check, the participant executes a 20° saccade towards a target on the right. Once the saccade is detected via a real-time algorithm, theflip command is given and a circular grating is displayed. The offset of the grating is triggered by the eyes crossing an invisible vertical boundary to the left of the target (white dotted line).**(B)**Overview of critical latencies, plotted relative to the moment that the saccade was detected online (time zero). Violin plots show median values and single-trial data points for saccade onset,flip latency, stimulus onset, stimulus offset, and saccade offset. Shading indicates the average duration that the grating was visible.**(C)**The viewer’s average horizontal gaze position.**(D)**Single-trial photodiode responses, sorted byflip latency in the trial (black line). Visible stripes result from the slight OLEDflicker during each refresh cycle.

In summary, using the OLED, it was possible to reliably turn the stimulus on and off again during a saccade with short delays. This made it possible to display the grating for two thirds of the movement’s duration, but not before or after. By using this setup together with simultaneous EEG recordings (Dimigen et al., 2011), we can assess whether and how the intrasaccadic stimulation of the retina (Campbell & Wurtz, 1978; Nicolas et al., 2021; Wurtz, 1969) contributes to eye movement-related neural activity in the EEG (Dimigen & Ehinger, 2021). These results are reported elsewhere.

## DISCUSSION

While the advantages of OLED panels in delivering excellent image quality are well recognized, these displays are not yet commonly used in studies on visual perception or cognitive neuroscience. In this study, we evaluated a promising model from a new generation of fast OLED gaming monitors; focusing on its temporal properties for monochromatic stimuli. We found that the 240 Hz monitor has excellent timing properties, consistent with the already promising results for much earlier OLED models (Cooper et al., 2013; Elze et al., 2013; Ito et al., 2013). Transition times andflip-to-90% latencies were significantly better than those of the tested LCDs and on par with those of a CRT monitor. Like a CRT monitor, the OLED was able to show true black, supporting high contrasts. Unlike a CRT monitor, however, the OLED was also able to display stimuli as continuous, almost square wave-like luminance patterns at their nominal duration (Elze, 2010b) and without slowly decaying offsets due to phosphor persistence. The monitor’s 240 Hz refresh rate supports fine-grained gradations in stimulus duration that are important for many paradigms in experimental psychology (Poth et al., 2018)^2^. Its fast transition times should also improve the measurement of visually-evoked brain potentials as compared to LCDs (Husain et al., 2009; Kaltwasser et al., 2009; Matsumoto et al., 2014; Nagy et al., 2011).

### Sources of luminance variation

We identified several factors that subtly influenced the luminance displayed by the OLED. During each 240 Hz refresh cycle, the monitor exhibited a very brief luminance dip (of up to 5%), aflicker that is typical for OLED panels. However, due to its high frequency, thisflicker should be imperceptible to the viewer and it only has a minor impact on average luminance over time.

We also assessed luminance variations due to screen location, viewing angle, and operating time. With a standard deviation between screen locations of 1.5%, the monitor’s luminance uniformity was decent and differences did not exceed 5.5% for any two screen locations. Whereas CRTs are barely affected by viewing angle (Brainard et al., 2002), poor viewing angles are a major drawback of LCDs. For the OLED, luminance decreased to 92% at viewing angle offsets of ±30°, the maximum range likely to be encountered in experiments with centralfixation ^3^. This value is at least in the range of those reported for CRTs (Ghodrati et al., 2015; their Fig. 3B), better than that obtained for the tested IPS panel (LCD-1; decrease to 80.5%) and significantly better than those reported in older tests of research-grade LCDs (Ghodrati et al., 2015; their Fig. 3B). Finally, we found that just like CRTs and LCDs (e.g., Brainard et al., 2002; Fox et al., 2014; Poth & Horstmann, 2017), the monitor needed to warm up for at least half an hour (e.g., 36 minutes to reach 99% of its maximum luminance) before any experiment.

### Controlling OLED-specific luminance artifacts

A major complication for precise experimentation with OLED panels is their inherent Auto Brightness Limiting feature, which regulates the panels brightness to balance power usage. Importantly, our measurements suggest that ABL can be prevented by operating the monitor at a moderate brightness setting of 40% which maps to a luminance of about 140 cd/m^2^. Given that many experiments are conducted in dimmed environments and that the monitor has excellent contrasts, this luminance may be sufficient for many use cases. As an alternative, we found that the monitor’s “Uniform Brightness” mode effectively prevented luminance changes due to ABL, allowing users to safely operate the monitor at a luminance of 260 cd/m^2^. Notably, this mode only worked under Windows, but not with an otherwise identical Linux setup. Overall, our results suggest that ABL-like saturation phenomena, already observed in earlier tests of OLEDs (Cooper et al., 2013; Elze et al., 2013; Ito et al., 2013), can be effectively prevented.

This was not true for another unexpected dimming phenomenon, which we called dynamic dimming. During the first 500 ms of their presentation, bright stimuli decreased in luminance by up to 5%. Interestingly, this decrease was seen exclusively for smaller stimuli subtending 10% or less of the screen. In contrast to ABL, this dimming occurred regardless of the monitor’s brightness setting and regardless of the Uniform Brightness mode. We are unaware of the technical mechanisms causing this seemingly unavoidable dimming. However, as mentioned, the magnitude of this luminance change was relatively small (about 5% for the smallest tested stimuli), which is less than, for example, the within-cycle luminance fluctuations observed with the tested LCDs (see Figure 1B). It may be tolerable for many experimental applications.

### Temporal properties support time-critical paradigms

We practically assessed the monitor’s temporal properties by implementing a fast flicker stimulation resembling that used in Rapid invisible frequency tagging (RIFT). Originally, RIFT was implemented with a 1440 Hz projector which can show 60 Hz sinusoidal flicker sequence at 24 frames/cycle (Minarik et al., 2023; Seijdel et al., 2023). Recently, it was demonstrated that attention-modulated RIFT signals can also be measured in EEG (Arora et al., 2024) at refresh rates of 480 Hz, yielding 8 frames per cycle. As shown here, the tested 240 Hz OLED can present 60 Hz flicker at four frames per cycle, allowing for the inclusion of two intermediate gray states per cycle to reduce visibility. Due to its short transition times, the OLED was able to properly display each luminance state for most of the 4.17 ms frame duration, a feat likely unachievable with an LCD. While a 240 Hz refresh rate is still insufficient for simultaneous attentional tracking at multiple neighboring frequencies (e.g., at 60 and 64 Hz; Arora et al., 2024), manufacturers have already announced gaming OLEDs capable of running at 480 Hz. Our results therefore suggest that even complex RIFT protocols should soon be feasible with consumer hardware.

Our second practical test was a gaze-contingent eye-tracking experiment that required the rapid presentation of a stimulus during a saccade, but not before or after. In our study, the time needed to detect that the eye was moving (*SaccOn*to*SaccDetect*interval) was 5.28 ms. This interval is independent of the monitor used and its exact duration also depends on the thresholds chosen to detect saccades in the offline analysis. Nevertheless, the low value of around 5 ms confirms the good performance of the online algorithm proposed by Schweitzer & Rolfs (2020)and suggests that this method is an alternative to classic “boundary techniques” (Rayner, 1975) where the saccade is only detected once the gaze exceeds a certain pixel limit.

More important in the present context was the display change latency, which depends critically on the monitor’s refresh rate, input lag, and rise time. Once the saccade was detected, the monitor was able to show the new stimulus at >90% luminance within 7.61 ms. For comparison, a monitor like the LCD-1, with its 144 Hz refresh rate, would have delayed this change by at least another 12 ms^4^. The value of 7.61 ms is significantly better than what we have previously achieved in our research with fast CRT monitors (e.g., 9.7 ms in Dimigen et al., 2012; 10.7 ms in Kornrumpf et al., 2016)^5^. It is also shorter than the latency reached with a high-speed projector by Schweitzer & Rolfs (2020)^6^.

Summing up the two intervals, the total latency from saccade onset to stimulus onset was below 13 ms. In practical terms, this meant that we could present the stimulus for the majority of the saccade duration, but not before or after, and also that fewer trials were lost due to late changes.

In principle, these good timing properties should also allow researchers to present multiple stimuli during a saccade, which could alleviate some methodological concerns about saccade-contingent paradigms. For example, to measure the preview benefit from parafoveal information, eye movement studies on reading often compare a condition with a correct presaccadic preview to those with an invalid preview. Because only the invalid conditions involve a change of the display, it is at least conceivable that this additional intrasaccadic transient contributes to the preview effects seen in fixation times (e.g., via oculomotor inhibition; Inhoff et al., 1998; O’Regan, 1990; Reingold & Stampe, 2000; Vitu-Thibault, 2007) or neural responses (Chase & Kalil, 1972; Dimigen et al., 2012; Michael & Stark, 1967). The tested OLED would be fast enough to insert an additional intrasaccadic transient (e.g., 1-2 frames of an x-letter mask) in all preview conditions (Buonocore et al., 2020), thereby matching them better in terms of low-level stimulation.

### Not tested: Motion, color, and longevity

Our tests focused on timing properties for monochromatic stimuli; motion and color were not examined. To reduce motion blur, it is generally advantageous to use a strobing display with a duty cycle that lasts only a fraction of the refresh cycle. This was the case for the older 60 Hz OLEDs tested for vision research (Elze et al., 2013; Matsumoto et al., 2014). In contrast, the model tested here is a “sample-and-hold” display, meaning that each image is displayed for the full frame duration (see Figure 1A). In many ways, this is a desirable property for a research display, since the stimulus durations actually match the nominal duration specified by the experimenter (Elze, 2010a). Although sample-and-hold displays result in more motion blur than strobing displays, the high refresh rate and fast transition times of the tested panel should largely mitigate this issue. Tests of color reproduction are found in recent product reviews (Baker, 2023; Simmons, 2023; Thauvette et al., 2024) and have reported a wide color gamut and good color accuracy as compared to LCDs.

Finally, we were not able to test the panel’s longevity. As mentioned, OLEDs are more vulnerable to thermal and electrical stress than other panels and degradation of their organic materials may cause long-term issues such as burn-in, color shifts, or a reduction in luminance. While several screen protection features must be disabled for precise stimulation, others, such as “pixel cleaning”, can be run in-between experiments. Whether or not there is a serious risk of burn-in with typical experimental usage at moderate brightness settings is presently unclear. In any case, it seems prudent to occasionally verify and recalibrate the monitor’s output.

### Conclusions

In our test, the latest generation of consumer-grade high-speed OLED gaming monitors demonstrated excellent temporal properties with short input lags, CRT-like transition times, and temporally independent and symmetric on- and offset responses. The self-lit pixels of the OLED support high contrasts and provide good viewing angles and luminance uniformity. With its 240 Hz refresh rate, the monitor seems suitable for paradigms in cognitive science requiring the fast presentation of stimuli with finely graded durations, as demonstrated here for a saccade-contingent paradigm. To prevent luminance artifacts, the monitor needs to be run at moderate brightness levels or within a special mode. Long-term tests are needed to assess the monitor’s longevity under laboratory conditions. Lastly, it is significant that the tested monitor costs only a fraction of dedicated LCDs and projector systems developed for vision science and EEG recordings. We hope that its strong performance can help to level the playing field by enabling researchers to implement time-critical paradigms regardless of their budget.

## OPEN PRACTICES STATEMENT

Data and Matlab code to reproduce all results and figures will be made available at https://osf.io/8h4fg/ upon publication. The study was not pre-registered.

## DECLARATIONS

### Ethics approval

According to the guidelines of the local ethics commission, the study was exempt from ethical approval.

### Consent to participate

The participant provided written informed consent.

### Consent for publication

All authors (O.D. and A.S.) consented to the manuscript being submitted for publication.

### Authors contributions

Conceptualization: OD. Methodology: OD, AS. Software: OD, AS. Investigation: OD, AS. Formal analysis: OD. Visualization: OD. Writing - Original draft: OD. Writing - Review & Editing: OD, AS.

## Footnotes

1 Despite the technique’s name, the full invisibility of theflicker has not been systematically confirmed. It may not hold under conditions where high-velocity (micro)saccades spread the flicker across the retina.

2 The resolution of possible stimulus durations could be even further enhanced by using the monitor together with the variable refresh rate feature of modern graphics cards, as proposed by (Poth et al., 2018). Because this approach requires that the stimulus lasts at least one frame and that its onset is predictable it is not feasible in tasks where the stimulation is triggered by behavior, as in our saccade task.

3 At a viewing distance of 50 cm, the left and right edges of the 27” OLED screen are located at ±30.5°.

4 This is the sum of a cycle duration (flip latency) that is 2.78 ms longer at 144 Hz plus a much longer flip-to-90% duration of 9.25 ms (see also Fig. 1B).

5 Some authors in reading research have reported shorter latencies with much slower monitors. We suspect that these measurements did not consider the time that the cathode ray needs to travel back to the vertical screen center, where sentences were presented.

6 State-of-the-art projectors can display preprogrammed stimulus sequences at 1440 Hz, but they can only react to real-time user behavior at the slower rate of 120 Hz in this mode (Schweitzer & Rolfs, 2020).

